# Flexible sensorimotor computations through rapid reconfiguration of cortical dynamics

**DOI:** 10.1101/261214

**Authors:** Evan D. Remington, Devika Narain, Eghbal A. Hosseini, Mehrdad Jazayeri

## Abstract

Sensorimotor computations can be flexibly adjusted according to internal states and contextual inputs. The mechanisms supporting this flexibility are not understood. Here, we tested the utility of a dynamical system perspective to approach this problem. In a dynamical system whose state is determined by interactions among neurons, computations can be rapidly and flexibly reconfigured by controlling the system‘s inputs and initial conditions. To investigate whether the brain employs such control strategies, we recorded from the dorsomedial frontal cortex (DMFC) of monkeys trained to measure time intervals and subsequently produce timed motor responses according to multiple context-specific stimulus-response rules. Analysis of the geometry of neural states revealed a control mechanism that relied on the system‘s inputs and initial conditions. A tonic input specified by the behavioral context adjusted firing rates throughout each trial, while the dynamics in the measurement epoch allowed the system to establish initial conditions for the ensuing production epoch. This initial condition in turn set the speed of neural dynamics in the production epoch allowing the animal to aim for the target interval. These results provide evidence that the language of dynamical systems can be used to parsimoniously link brain activity to sensorimotor computations.

## Introduction

Humans and nonhuman primates are capable of generating a vast array of behaviors, a feat dependent on the brain‘s ability to produce a vast repertoire of neural activity patterns. However, identifying the mechanisms by which the brain flexibly selects neural activity patterns across a multitude of contexts remains a fundamental and outstanding problem in systems neuroscience.

Here, we aimed to answer this question using a dynamical systems approach. Work in the motor system has provided support for a hypothesis that movement-related activity in motor cortex can be described at the level of neural populations and viewed as low dimensional neural trajectories of a dynamical system (Churchland et al. 2010; Churchland et al. 2012; Seely et al. 2016; Fetz 1992; Michaels et al. 2016). More recently, a dynamical systems view has been used to provide explanations for neural trajectories in premotor and prefrontal cortical areas in various cognitive tasks (Mante et al. 2013; Rigotti et al. 2010; Carnevale et al. 2015; Hennequin et al. 2014; Rajan et al. 2016). This line of investigation has been complemented by efforts in developing, training, and analyzing recurrent neural network models that can emulate a range of motor and cognitive behaviors, leading to novel insights into the underlying latent dynamics (Mante et al. 2013; Hennequin et al. 2014; Sussillo et al. 2015; Chaisangmongkon et al. 2017; Wang et al. 2017). These early successes hold promise for the development of a more ambitious “computation-through-dynamics” (CTD) as a general framework for understanding how activity patterns in the brain support flexible behaviorally-relevant computations.

The behavior of a dynamical system can be described in terms of three components: (1) the interaction between state variables that characterize the system's latent dynamics, (2) the system‘s initial state, and (3) the external inputs to the system. Accordingly, the hope for using the mathematics of dynamical systems to understand flexible generation of neural activity patterns and behavior depends on our ability to understand the co-evolution of behavioral and neural states in terms of these three components. Assuming that synaptic couplings between neurons and other biophysical properties are approximately constant on short timescales (i.e. trial to trial), we asked whether behavioral flexibility can be understood in terms of adjustments to initial state and external inputs.

There is evidence that certain aspects of behavioral flexibility can be understood through these mechanisms. For example, it has been proposed that preparatory activity prior to movement initializes the system such that ensuing movement-related activity follows the appropriate trajectory (Churchland et al. 2010). Similarly, the presence of a context input can enable a recurrent neural network to perform flexible rule-(Mante et al. 2013; Song et al. 2016) and category-based decisions (Chaisangmongkon et al. 2017). However, whether these initial insights would apply more broadly and generalize when both inputs and initial conditions change is an important outstanding question.

For many behaviors, distinguishing the effects of the synaptic coupling, inputs and initial conditions in neural activity patterns is challenging. For example, neural activity during a reaching movement is likely governed by both local recurrent interactions and distal inputs from time-varying and condition-dependent reafferent signals (Todorov & Jordan 2002; Scott 2004; Pruszynski et al. 2011). Similarly, in many perceptual decision making tasks, it is not straightforward to disambiguate the sensory drive from recurrent activity representing the formation of a decision and the subsequent motor plan (Mante et al. 2013; Meister et al. 2013; Thura & Cisek 2014). This makes it difficult to tease apart the contribution of recurrent dynamics governed by initial conditions from the contribution of dynamic inputs (Sussillo et al. 2016). To address this challenge, we designed a sensorimotor task for nonhuman primates in which animals had to measure and produce time intervals using internally-generated patterns of neural activity in the absence of potentially confounding time varying sensory and reafferent inputs. Using a novel analysis of the geometry and dynamics of *in-vivo* activity in the dorsal medial frontal cortex (DMFC) and *in-silico* activity in recurrent neural network models trained to perform the same task, we found that behavioral flexibility is mediated by the complementary action of inputs and initial conditions controlling the structural organization of neural trajectories.

## Results

### Ready, Set, Go (RSG) task

Our aim was to ask whether flexible control of internally-generated dynamics could be understood in terms of systematic adjustments made to initial conditions and external inputs of a dynamical system. We designed a “Ready, Set, Go” (RSG) timing task to directly investigate the role of these two factors. The basic sensory and motor events in the task were as follows: following fixation of a central spot, monkeys viewed two peripheral visual flashes (“Ready” followed by “Set”) separated by a sample interval, *t s*, and produced an interval, *t p*, after Set by making a saccade to a visual target that was presented throughout the trial. In order to obtain juice reward, animals had to generate *t* p as close as possible to a target interval, *t t* (Figure 1B), which was equal to *t s* times a “gain factor”*g* (*t t* = *gt s*). The demand for flexibility was imposed in two ways (Figure 1C). First, *t s* varied between 0.5 and 1 sec on a trial-by-trial basis (drawn from a discrete uniform “prior” distribution). Second, *g* switched between 1 (*g* =1 context) and 1.5 (*g* =1.5 context) across blocks of trials)Figure 1D, mean block length = 101, std = 49 trials).

To verify that animals learned the task (Figure 1E), we used regression analyses to assess the dependence of *t p* on *t s* and *g*. First, we analyzed the relationship between *t s* and *t p* within each context (*t p* =β0 + β 1 *t s*). Results indicated that *t p* increased monotonically with *t s* for both contexts (β 1 > 0, p << 0.001 for all sessions). Next, we assessed the influence of gain on *t p* in several complementary analyses. First, we compared regression slopes relating *t p* to *t s* within each context. The slopes were significantly higher in the *g* =1.5 compared to *g* =1 context (meanβ 1 = 0.84 vs. 1.2; signed-rank test p = 0.002, n = 10 sessions; Figure 1E, inset). Second, we fit a regression model to behavior across both gains that included additional regressors for gain and its interaction with *t s* (*t p* =β 0 +β 1 *t s* +β 2 *g* +β3 *gt s*). Results indicated a significant positive interaction between *t s* and *g* (meanβ 3 =, to the number of trials following a context switch to determine how fast monkeys adjusted their behavior. There was no evidence for a slow adaptation of *t p* as a function of number of trials after switch (one-tailed test forβ 0.73;β_3_ > 0, p < 0.0001 in each session). Finally, we fit a regression model relating *t p*, z-scored for each *t s* 1 in first 25 trials after switch, p > 0.25), indicating that the switching was rapid. Together, these results confirmed that animals used an estimate of *t s* to compute *t p* and flexibly adjusted their responses according to the gain information.

For both gains, responses were variable, and average responses exhibited a regression to the mean (meanβ1 < 1, p = 0.005 for *g* =1, and mean β 1.5, p = 0.0001 for *g* =1.5, one-sided signed-rank test). As with previous work (Jazayeri & Shadlen 2015; Acerbi et al. 2012; Miyazaki et al. 2005; Jazayeri & Shadlen 2010), behavior was accurately captured by a Bayesian model (Figure 1E, Methods) indicating that animals integrated their knowledge about the prior distribution, the sample interval and the gain to optimize their behavior.

**Figure 1.**
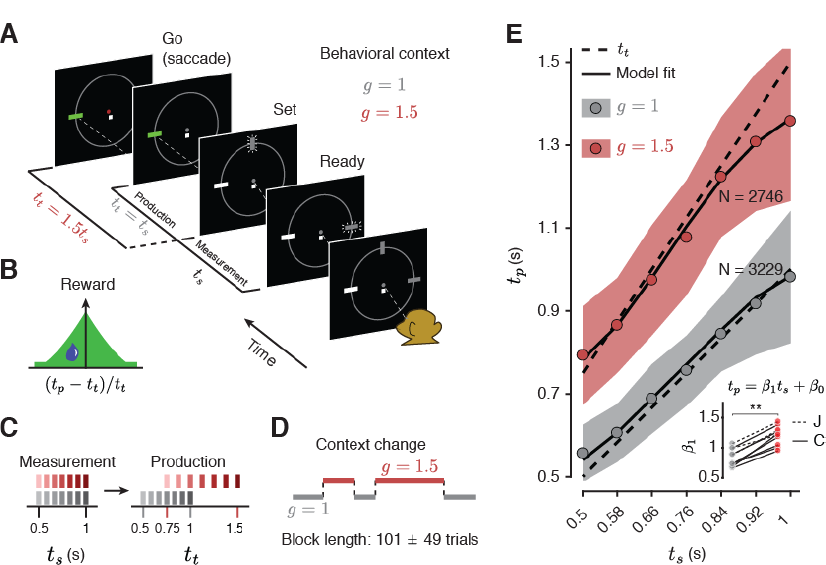
The RSG task and behavior. (**A**) RSG task. On each trial, three rectangular stimuli termed “Ready,” “Set,” and “Go” were shown on the screen arranged in a semi-circle. Following fixation, Ready and Set were extinguished. After a random delay, first Ready and then Set stimuli were flashed (small lines around the rectangles signify flashed stimuli). The time interval between Ready and Set demarcated a sample interval, *t* _*s*_. The monkey‘s task was to generate a saccade (“Go”) to a visual target such that the interval between Set and Go (produced interval, *t* _*p*_) was equal to a target interval, *t* _*t*_, of *t* _*s*_ multiplied by a gain factor, *g*. The animal had to perform the task in two behavioral contexts, one in which *t* _*t*_ was equal to *t* _*s*_ (*g* =1 context), and one in which *t* _*t*_ was 50% longer than *t* _*s*_ (*g* =1.5 context). The context was cued by the color of fixation and the position of a context stimulus (small white square below the fixation) throughout the trial. (B) Animals received juice reward when the error between *t* _*p*_ and *t* _*t*_ was small, and the reward magnitude decreased with the the size of error (see Methods for details). On rewarded trials, the saccadic target turned green (panel A). (C) For both contexts, *t* _*s*_ was drawn from a discrete uniform distribution with seven values equally spaced from 0.5 to 1 sec (left). The values of *t* _*s*_ were chosen such that the corresponding values of *t* _*t*_ across the two contexts were different but partially overlapping (right). (D) The context changed across blocks of trials. The number of trials in a block was varied pseudorandomly (mean and std shown). (E) *t* _*p*_ as a function of *t* _*s*_ for each context across all recording sessions. Circles indicate mean *t* _*p*_ across all sessions, shaded regions indicate +/− one standard deviation from the mean, dashed lines indicate *t* _*t*_, and solid lines are the fits of a Bayesian observer model to behavior. Inset: Slope of the regression line (β 1) relating *t* _*p*_ to *t* _*s*_ in the two contexts. Regression slopes were larger in the *g* =1.5 context, with a significant interaction between *t* _*s*_ and *g* (p < 0.0001) for all sessions (see text; ** indicates p < 0.002 for signed-rank test). In all panels, different shades of gray and red are associated with *g* =1 and *g* =1.5, respectively.

### Neural activity in the RSG task

To assess the neural computations in RSG, we focused on the dorsal region of the medial frontal cortex (DMFC) comprising supplementary eye fields, supplementary motor area and presupplementary motor area. DMFC is a natural candidate for our task because it plays a crucial role in timing as shown by numerous studies in humans (Halsband et al. 1993; Rao et al. 2001; Coull et al. 2004; Pfeuty et al. 2005; Macar et al. 2006; Cui et al. 2009), monkeys (Okano & Tanji 1987; Merchant et al. 2013; Kunimatsu & Tanaka 2012; Isoda & Tanji 2003; Romo & Schultz 1992; Merchant et al. 2011; Mita et al. 2009; Ohmae et al. 2008; Kurata & Wise 1988), and rodents (Matell et al. 2003; Kim et al. 2009; Smith et al. 2010; Kim et al. 2013; Xu et al. 2014; Murakami et al. 2014), and because it is involved in context-specific control of actions (Isoda & Hikosaka 2007; Ray & Heinen 2015; Yang & Heinen 2014; Shima et al. 1996; Matsuzaka & Tanji 1996; Brass & von Cramon 2002).

We recorded from 326 units (127 from monkey C and 199 from monkey J) in DMFC. Between 11 and 82 units were recorded simultaneously in a given session, however in this study, we combined data across all units irrespective of whether they were recorded simultaneously. Firing patterns were heterogeneous and varied across units, task epochs, and experimental contexts. In the Ready-Set epoch, responses were modulated by both gain and elapsed time (e.g. units #1, 3, and 5, Figure 2A). For many units, firing rate modulations underwent a salient change at the earliest expected time of Set (0.5 sec). For example, responses of some units increased monotonically in the first 0.5 sec but decreased afterwards (Figure 2A, units #1, 3).

Following Set, firing rates were characterized by a mixture of 1) transient changes after Set (unit #1 and 3), 2) sustained modulations during the Set-Go epoch (units #1 and 5), and 3) monotonic changes in anticipation of the saccade (units #1, 2 and 4). These characteristics were not purely sensory or motor and varied systematically with *t s* and gain. For example, the amplitude of the early transient response (unit #1) depended on both *t s* and gain, indicating that it was not a visually-triggered response to Set. The same was true for the sustained modulations after Set and activity modulations prior to saccade initiation.

We also examined the representation of *t s* and gain across the population by projecting the data on dimensions along which activity was strongly modulated by context and interval in state-space (i.e. the space spanned by the firing rates of all 326 units; see Methods). Similar to individual units, population activity was modulated by both elapsed time and gain during both the Ready-Set (Figure 2B) and Set-Go (Figure 2C) epochs. We used this rich dataset to investigate whether the flexible adjustment of intrinsic dynamics across the population with respect to *t s* and gain could be understood using the language of dynamical systems.

**Figure 2.**
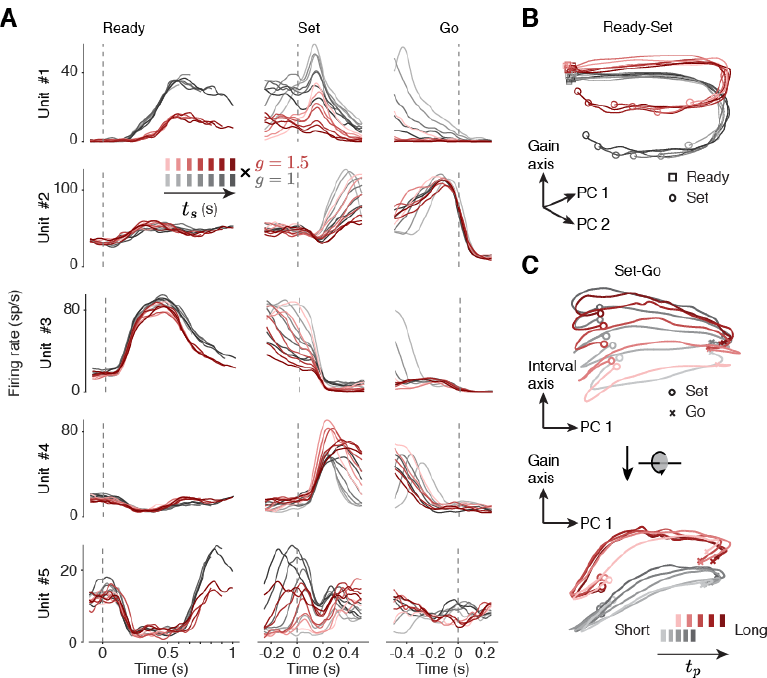
Neural responses in dorsomedial frontal cortex (DMFC) during the RSG task. (**A**) Firing rates of 5 example units during the various phases of the task aligned to Ready (left column), Set (middle) and Go (right). Responses aligned to Ready and Set were sorted by *t* _*s*_. Responses aligned to Go were sorted into 5 bins, each with the same number of trials, ordered by *t* _*p*_. Gray and red lines correspond to activity during the *g* =1 and *g* =1.5 contexts, respectively, with darker lines corresponding to longer intervals. (B) Visualization of population activity in the Ready-Set epoch sorted by *t* _*s*_. The “gain axis” corresponds to the axis along which responses were maximally separated with respect to context. The other two dimensions (“PC 1 & PC 2”) correspond to the first two principal components of the data after removing the context dimension. (C) Visualization of population activity in the Set-Go epoch sorted into 5 bins, each with the same number of trials, ordered by *t* _*p*_. Top: Activity plotted in 2 dimensions spanned by PC 1 and the dimension of maximum variance with respect to *t* _*p*_ within each context (“Interval axis”). Bottom: Same as Top rotated 90 degree (circular arrow) to visualize activity in the plane spanned by the context axis (“Gain axis”) and PC 1. In both panels, PC1 was computed after removing the variance explained along the Interval axis and Gain axis dimensions. Squares, circles, and crosses in the state space plots represent Ready, Set, and Go, respectively.

### Flexible neural computations: a dynamical systems perspective

We pursued the idea that neural computations responsible for flexible control of saccade initiation time can be understood in terms of the behavior of a dynamical system established by interactions among neurons. To formulate a rigorous hypothesis for how a dynamical system could confer such flexibility, we considered the goal of the task and worked backwards logically. The goal of the animal is to flexibly control the saccade initiation time to a fixed target. Previous motor timing studies proposed that saccade initiation is triggered when the activity of a subpopulation of neurons with monotonically increasing firing rates (i.e., “ramping“) reaches a threshold (Mita et al. 2009; Kunimatsu & Tanaka 2012; Romo & Schultz 1987; Roitman & Shadlen 2002; Hanes & Schall 1996; Tanaka 2005; Maimon & Assad 2006). For these neurons, flexibility requires that the slope of the ramping activity be adjusted (Jazayeri & Shadlen 2015). More recently, it was found that actions are initiated when the collective activity of neurons with both ramping and more complex activity patterns reach an action-triggering state (Churchland et al. 2006; Wang et al. 2017), and that flexible control of initiation time can be understood in terms of the speed with which neural activity evolves toward that terminal state (Wang et al. 2017).

In a dynamical system, the speed with which activity evolves over time is determined by the derivative of the state. If we denote the state of the system by, the derivative is usually specified by two factors, a function of the current state, *f*(*X*), and an external input, *U*, that may be constant or context- and/or time-dependent:

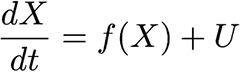

When analyzing the collective activity of a specific population of neurons, this formulation has a straightforward interpretation. The state represents the collective firing rate of neurons under investigation, accounts for the interactions among those neurons, and corresponds to external input from another population of neurons, possibly controlled by an external sensory drive. The only additional information needed to determine the behavior of this system is its initial condition, *X*_0_, which specifies the initial neural state prior to generating a desired dynamic pattern of activity.

To assess the utility of the dynamical systems perspective for understanding behavioral flexibility, we assumed that (i.e., synaptic coupling in DMFC) is fixed across trials. This leaves inputs and initial conditions as the only “dials” for achieving flexibility (Figure 3). To formalize a set of concrete hypotheses for the potential role of inputs and initial conditions, we first focused on behavioral flexibility with respect to *t s* for each gain context. How can a dynamical system adjust the speed at which activity during Set-Go evolves in a *t*_*s*_-dependent manner? In RSG, within each context, there are no sensory inputs (exafferent or reafferent) that could serve as a *t*_*s*_-dependent input drive. Therefore, we hypothesized that the *t*_*s*_-dependent adjustment of speed in the Set-Go epoch results from a parametric control of initial conditions at the time of Set. The corollary to this hypothesis is that the time-varying activity during the Ready-Set epoch is responsible for adjusting this initial condition based on the desired speed during the ensuing Set-Go epoch (Wang et al. 2017).

Second, we asked how speed might be controlled across the two gain contexts. One possibility is to establish initial conditions that generalize across the two contexts (Figure 3A). To do so, initial conditions must vary with, which has implicit information about both gain and *t s* (i.e.,. If both gain and *t s* are encoded by initial conditions, we would expect neural trajectories to form a single organized structure with respect to the target time (*t t* = *gt s* speed requirements associated with producing *t t* = *gt s*. In the extreme case, neural trajectories associated with the same value of *gt s* across the two contexts (e.g, 1.5×0.5 and 1.0×0.75) should terminate in the same state at the time of Set and should evolve along identical trajectories during the Set-Go epoch. We refer to this solution as *A 1* (Figure 3A).

Alternatively, DMFC responses may rely on a persistent gain-dependent input to adjust speed across the two gain contexts (Figure 3B). As exemplified by recurrent neural network models, in dynamical systems, a persistent input can rapidly reconfigure computations by driving the system to different regions of the state space (Mante et al. 2013; Sussillo et al. 2015; Hennequin et al. 2014; Chaisangmongkon et al. 2017; Song et al. 2016). This solution, which we refer to as *A 2*, predicts a qualitatively different geometrical organization of, with two key features. First, there should be a gain-dependent organization forming two sets of neural trajectories in two different regions of the state space. Second, neural trajectories should be organized with respect to *t s* and *t p* (i.e., within each context) but not necessarily with respect to *t t* (i.e., across contexts). Because the context information in RSG was provided as an external visual input (fixation cue), and was available throughout the trial, we predict that this solution offers the more plausible prediction for how the brain might solve the task.

Therefore, the dynamical systems perspective in RSG leads us to the following specific hypotheses: 1) the evolution of activity in the Ready-Set epoch parametrizes the initial conditions needed to control the speed of dynamics in the production epoch for each context, and 2) the context cue acts as a persistent external input leading the system to establish structurally similar yet distinct sets of neural trajectories associated with the two gains, and no *t p*,related structure across contexts, consistent with A 2.

Visualization of neural trajectories from Set to Go in state space (Figure 3C, same as in Figure 2C) provided qualitative support for these hypotheses. First, within each context, neural trajectories for different *t p* bins were clearly associated with different initial conditions and remained separate and ordered throughout the Set-Go epoch. Second, context information seemed to displace the entire group of neural trajectories to a different region of neural state space without altering their relative organization as a function of *t p*. Third, indexing time along nearby trajectories suggested that the speed with which responses evolved along each trajectory was systematically related to the desired *t t*; i.e, slower for longer *t t*. To validate these observations quantitatively, we neural trajectories compared to *A 1 A 2*. Figure 3C) provided developed an analysis technique which we termed “ki nematic analysis of ne ural t rajectories” (KiNeT) that helped us measure the relative speed and position of multiple, possibly curved (Figure S1), neural trajectories.

**Figure 3.**
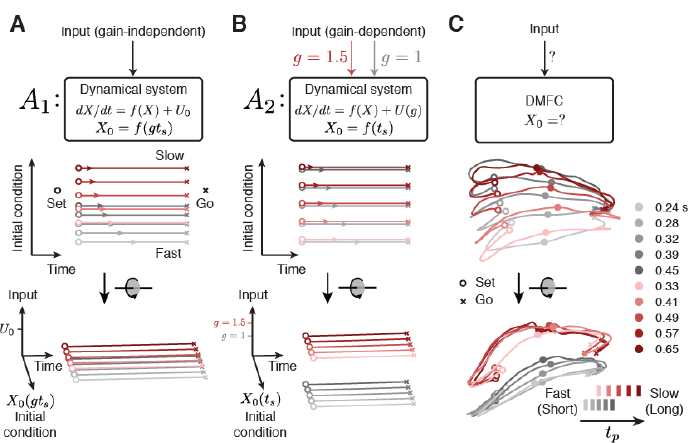
Dynamical systems predictions for the RSG task. (**A,B**) Schematic illustrations for dynamical systems solutions to generalize RSG across contexts through manipulation of initial conditions or external inputs. (A) Gain-control by initial condition (*A* _*1*_). Top: The target interval *t* _*t*_= *gt* _*s*_ (*g*, gain, *t* _*s*_, sample interval) is encoded by the initial conditions (*X* _*0*_(*gt* _*s*_)) generated during the Ready-Set epoch (not shown). Middle: After the Set cue (open circles), activity evolves towards an action-triggering state (crosses) with a speed (colored arrows) fully determined by position along the initial condition subspace (ordinate). Activity across contexts is organized according to *t* _*t*_= *gt* _*s*_. Bottom: same trajectories, rotated to show an oblique view. Trajectories are separated only along the initial condition axis across both contexts such that trajectory structure reflects *t* _*t*_ explicitly. There is no separation along the Input axis. (B). Gain-control by external input (*A* _*2*_). Top: *t* _*s*_ is encoded by initial conditions (*X* _*0*_(*t* _*s*_)), and a persistent context-dependent input encodes the gain (red and gray arrow for the two gains). Middle: within each context, trajectories associated with the same *t* _*s*_ evolve along the same position on the initial condition axis at different speeds due to the context-dependent input. Activity is organized according to *t* _*s*_ and not *t* _*t*_. Bottom: oblique view. A context-dependent external input creates two sets of neural trajectories in the state space for the two contexts in the Set-Go epoch. This input controls speed in conjunction with *t* _*s*_-dependent initial conditions, generating a structure which reflects *t* _*s*_ and *g* explicitly, but not *t* _*t*_. In both *A* _*1*_ and *A* _*2*_, responses would be initiated when activity projected onto the time axis reaches a threshold. (C) DMFC data. Top: unknown mechanism of RSG control in DMFC. Middle, bottom: 3-dimensional projection of DMFC activity in the Set-Go epoch (from Figure 2C). Middle: qualitative assessment indicated that neural trajectories within each context for different *t* _*p*_ bins were associated with different initial conditions and remained separate and ordered through the response. Bottom: Across the two contexts, neural trajectories formed two separated sets of neural trajectories without altering their relative organization as a function of *t* _*p*_. Both of these features were consistent with *A* _*2*_. Filled circles depict states along each trajectory at a constant fraction of the trajectory length, illustrating speed differences across trajectories.

### Control of neural trajectories by initial condition within contexts

We first employed KiNeT to validate that animals’ behavior was predicted by the speed with which neural trajectories evolved over time. We reasoned that neural states evolving faster will reach the same destination on the trajectory in a shorter amount of time. Therefore, we estimated relative speed across the trajectories by performing a time alignment to identify the times when neural activity reached nearby points on each trajectory (Figure 4A). We then used this approach to analyze the geometrical structure of trajectories through the Set-Go epoch.

To perform KiNeT, we binned trials from each gain and recording session into five groups according to *t*_*p*_. Neural responses from these trials were averaged, then PCA was applied to generate five neural trajectories within the state space spanned by the first 10 PCs that explained 89% of variance. We denote each trajectory by Ω [*i*](*t*) (or Ω [*i*] for shorthand; a table with definitions of all symbols is provided in Methods) where indexes the trajectory and represents elapsed time since Set. We estimated speed and position along each Ω [*i*] relative to the trajectory associated with the middle (third) bin, which we refer to as the reference trajectory Ω[ref]. We denoted neural states on the reference trajectory by s[ref][j], where j indexes states through time along Ω[ref]. We used curly brackets to refer to a collection of indices. For example,s[ref]{j} refers to all states on Ω[ref], and corresponds to the time points on Ω[ref] associated with those states.

For each *s*[ref][*j*], we found the nearest point on all non-reference trajectories (*i*≠) as measured by Euclidean distance. We denoted the collection of the nearest states onΩ[] by*s*[*[i]{j}*, and the corresponding time points by*t*[*i*{*j*}. The corresponding time points along different trajectories provided the means for comparing speed: if *t*[*i*]{/,i>j}were systematically greater than *t*[ref]{*j*}, we could conclude thatΩ[*i*] evolves at a slower speed compared toΩ[ref] (Figure 4A). This relationship can be readily inferred from the slope of the line that relates *t*[*i*{*j*} to*t*[ref]{*j*}. While a unity slope indicates that the speeds are the same, higher and lower values would indicate slower and faster speeds of Ω [*i*] compared to Ω[ref], respectively.

Applying KiNeT to neural trajectories in the Set-Go epoch indicated that Ω{*i*} evolved at similar speeds immediately following the Set cue (unity slope). Later, speed profiles diverged such that neural trajectories associated with longer intervals slowed down and and trajectories associated with shorter intervals sped up for both gain contexts (Figure 4B). This is consistent with previous work that the key variable predicting *t p* is the speed with which neural trajectories evolve (Wang et al. 2017). One common concern in this type of analysis is that averaging firing rates across trials of slightly different duration could lead to a biased estimate of neural trajectory. To ensure that our estimates of average speed were robust, we applied KiNeT to neural trajectories while aligning trials to Go instead of Set. Results remained unchanged and confirmed that the speed of neural trajectories predicted *t p* across trials (Figure S2).

Having validated speed as the key variable for predicting *t p*, we focused on our first hypothesis that the evolution of activity in the Ready-Set epoch parametrizes the initial conditions needed to control the speed of dynamics in the production epoch for each context. Because speed is a scalar variable and has an orderly relationship to *t p*, this hypothesis predicts that the neural trajectories (and their initial conditions) should also have an orderly organizational structure with respect to *t p*. In other words, there should be a systematic relationship between the vectors connecting nearest points across neural trajectories and the *t p* to which they correspond. We tested this prediction in two complementary ways. First, we performed an *analysis of direction* testing whether the vectors connecting nearby trajectories were more aligned than expected by chance. Second, we performed an *analysis of distance* asking whether the distance between the reference trajectory and the other trajectories respected the distance between the corresponding speeds.

<u>Analysis</u> <u>of</u> <u>direction</u>. We used KiNeT to measure the angle between vectors connecting nearest points (Euclidean distance) across consecutive trajectories ordered by *t p*. Let us use 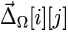 to denote the difference vector (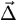) connecting nearest points across trajectories (subscript Ω) between *s*[*i*][*j*] and *s*[*i*+1][*j*]. According to our hypothesis, the direction of 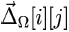 should be similar to 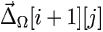 connecting to *s*[*i*+1]{*j*]. To test this, we measured the angle between these two difference vectors, denoted byθΩ[*i*][*j*]. The null hypothesis of unordered trajectories predicts that 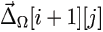 and 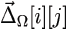 should be unaligned on average (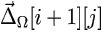 = 90 degrees; bar signifies mean of the angles over the index in curly brackets). Results indicated that θΩ[*i*][*j*] was substantially smaller than 90 degrees for both contexts (Figure 4C). This provides the first line of quantitative evidence for an orderly organization of neural trajectories with respect to *t p*.

Analysis of distance

We used KiNeT to measure the length of the vectors connecting nearest points on Ω[*i*]and Ω[ref], denoted by *D*[*i*][*j*], at different time points ([*j*). This analysis revealed that trajectories evolving faster than Ω{ref]and those evolving slower than Ω[ref] were located on the opposite sides of, and that the magnitude *D*[*i*[*j*] of increased progressively for larger speed differences (Figure 4D). This analysis provided clear evidence that, for each context, the relative position of neural trajectories and their initial conditions in the state space were predictive of *t p*.

To further substantiate the link between the geometry of neural trajectories and behavior, we asked whether trial-by-trial fluctuations of *t p* for each *t s* could be explained in terms of systematic fluctuations of speed and location of neural trajectories in the state space. We reasoned that fluctuations of *t p* partially reflect animals’ misestimation of *t s*. This predicts that larger values of *t p* for the same *t s* result from slower neural trajectories whose location in state space are biased toward longer values of *t s*. We tested this prediction by using KiNet to examine the relative geometrical organization of neural trajectories associated with larger and smaller values of *t p* for the same *t s*. Results indicated that neural trajectories that correspond to larger values of *t* slower speeds and were shifted in state space toward larger values of *ts* (*p* (Figure S3). This analysis extends the correspondence between behavior and the organization of neural trajectories to include animals’ trial-by-trial variability. Together, these results provide strong evidence for our first hypothesis: that activity during Ready-Set epoch parametrically adjusts the system‘s initial condition (i.e., neural state at the time of Set), which in turn controls the speed of neural trajectory in the Set-Go epoch and the consequent *t*

**Figure 4.**
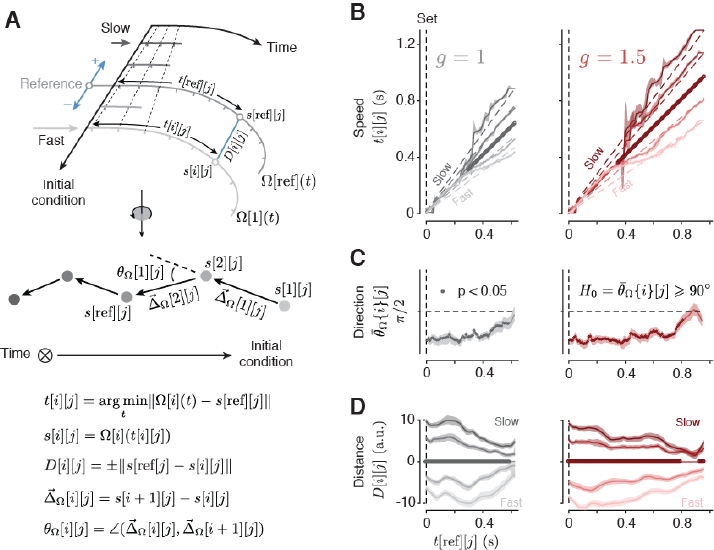
Kinematic analysis of neural trajectories (KiNeT). (A) Illustration of KiNeT. Top: a collection of trajectories Ω {*i*} originate from Set, organized by initial condition, and terminate at Go. Tick marks on the trajectories indicate unit time. Darker trajectories evolve at a lower speed as demonstrated by the distance between tick marks and the dashed line connecting tick marks. KiNeT quantifies the position of trajectories and the speed with which states evolve along them relative to a reference trajectory (middle trajectory, Ω[ref]).To do so, it finds a collection of states*s*[*i*]{*j*} on each Ω[*i* that are closest to Ω [ref] through time. Trajectories which evolve at a slower speed require more time to reach those states leading to larger values of *t*[*i*][*j*]. KiNet quantifies relative position by a distance measure, *D*[*i*][*j*](distance between Ω[*i*]and Ω[ref] at*t*[*i*][*j*]) that is signed (blue arrows) and is considered positive when Ω[*i*] corresponds to larger values of *t* _*p*_ (slower trajectories). Middle: trajectories rotated such that the time axis is normal to the plane of illustration, denoted by a circle with an inscribed cross. Filled circles represent the states aligned to for a particular *j*. Vectors 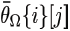 connect states on trajectories of shorter to longer *t* _*p*_. Angles θ_Ω_[*i*][*j*] between successive 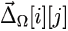 provide a measure of *t* _*p*_-related structure. Bottom: equations defining the relevant variables. (B). Speed of neural trajectories compared to computed for each context separately. Shortly after Set, all trajectories evolved with similar speed (unity slope). Afterwards, associated with shorter *t* _*s*_ evolved faster than as indicated by a slope of less than unity (i.e., smaller than), associated with longer *t* _*s*_ evolved slower than Ω[ref]. Filled circles on the unity line indicate*j* values for which *t*[*i*][*j*] was significantly correlated with *t*^p^[*i*][*j*] (bootstrap test, r > 0, p < 0.05, n = 100). (C). Relative position of adjacent neural trajectories computed for each context separately. 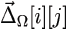 (bar signifies average across trajectories) were significantly smaller than 90 degrees (filled circle) for the majority of the Set-Go epoch (bootstrap test, 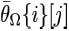 < 90, p < 0.05, n = 100) indicating that 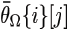 were similar across Ω[*i*]. (D) Distance of neural trajectories to Ω[ref] computed for each context separately. Distance measures (*D*[*i*][*j*]) indicated that Ω{*i*} had the same ordering as *t*^p^{*i*}. Significance tested using bootstrap samples for each *j* (p < 0.05, n = 100).

### Control of neural trajectories across contexts by external input

To identify the mechanism by which flexible speed control might be generalized across contexts, we first tested whether both gain and *t s* are encoded by initial conditions (*A 1*). According to this alternative, neural trajectories should follow the organization of *t p* across both contexts (Figure 3A), in addition to within each context (Figure 4C). To test *A 1*, we sorted neural trajectories across the two contexts according to *t p* (Figure 5A, top), and asked whether the angle between vectors connecting nearest points (θΩ{*i*][*j*]) was significantly less than 90degrees (Figure 5A, bottom). Unlike the within-context results (Figure 4C), when neural trajectories from both contexts were combined, the angle between nearby neural trajectories was significantly larger than 90 degrees (p < 0.05 for all *j*; Figure 5B). This indicates that trajectories across contexts do not have a orderly relationship to *t t* (*A 1*: less than 90 deg) even though they exhibit a structural organization that deviates from randomness (90 deg).

Next, we investigated the hypothesis that the context cue acts as a persistent external input (*A 2* Figure3B), leading the system to establish structurally similar but distinct collections of neural trajectories across contexts (Figure 6A,B). This hypothesis can be broken down to a set of specific geometrical constraints in the Set-Go epoch. We determined whether the data met these constraints by testing whether the converse of each could be rejected, as illustrated in Figure 6C-F. If we denote the collection of neural trajectories in the two contexts by Ω^g^=1{*i*}, and Ω=1.5{*i*}, these constraints and tests can be formalized as follows:

1. Ω^g^=1{*i*} and Ω=1.5{*i*} should evolve in the same direction as a function of time with different average speeds (i.e. slower for). If the converse were true (i.e., trajectories evolving in different directions,Figure 6C, left), we would expect no systematic relationship between time points across the two contexts. Results from KiNeT across contexts (see Methods) revealed a monotonically increasing relationship between *t*^g^=1[ref]{*j*} and *t*^g^=1.5[ref]{*i*}, confirming that Set-Go trajectories across contexts evolved in the same direction (Figure 6C, right). Moreover,*t*^g^=1[ref]{*j*} had a higher rate of change than indicating that average speeds were slower in the g=1.5 condition. This suggests that speed control played a consistent role across contexts (Figure 6A).
2. Ω^g^=1{*i*} and Ω^g^=1.5{*i*}should be organized similarly with respect to *t p*. In other words, the vector that connects nearby points in Ω^g^=1{*i*} should be aligned to its counterpart that connect nearby points in Ω^g^=1.5{*i*}. To evaluate this constraint, we used the angle between pairs of vectors that connect nearby points within each context. We use an example to illustrate the procedure (Figure 6B). Consider one vector connecting nearby points in two successive neural trajectories in the gain of 1 (e.g Ω^g^=1[1].and Ω^g^=1[2]), and another vector connecting the corresponding points in the gain of 1.5 (e.g., Ω^g^=1.5[1] and Ω^g^=1.5[2]). A similar orientation between the two groups of trajectories (Figure 6A) would cause the angle between these vectors (θ^g^=1[1]) to be significantly smaller than 90 degrees. If instead, Ω^g^=1{*i*}and Ω^g^=1.5{*i*} were oriented differently (Figure 6D, left) or had no consistent relationship, these vectors would be on average orthogonal. Using KiNeT, we found that this angle (θ^g^[*i*][*j*]) was consistently smaller than 90 degrees throughout the Set-Go epoch, providing quantitative evidence that the collection of neural trajectories associated with the two gains were structurally similar (Figure 6A).
3. If context information is provided as a tonic input, Ω^g^=1{*i*} and Ω^g^=1.5{*i*} should be separated in state space along a context axis throughout the Set-Go epoch. To verify this constraint, we assumed that neural trajectories for each context were embedded in distinct manifolds and compared the minimum distance between the two manifolds () to an analogous distance metric within each manifold (Figure 6B; see Methods). These distance measures should be the same if the groups of trajectories associated with the two contexts overlap in state space (Figure 6E, left). However, we found distances to be substantially larger across contexts compared to within contexts(Figure 6E, right). This confirms that the groups of trajectories associated with the two contexts were separated in state space (Figure 6A).
4. The results so far reject a number of alternative hypotheses (Figure 6C,D,E) and leave out two possibilities: either Ω^g^=1{*i*} and Ω^g^=1.5{*i*}are separated along the same dimension that separates trajectories within each context (Figure 6F, left), or they are separated along a distinct input axis in accordance with *A 2* (Figure 6A). To distinguish between these two, we asked whether the vector associated with the minimum distance (*D*^g^[*j*]) 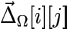 was aligned to vectors connecting nearby states within each context (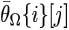). Analysis of the angle between these vectors (θ^g^,Ω[*j*]) indicated that the two were orthogonal for almost all *j* (Figure 6F,right). This ruled out the remaining possibility that trajectories across contexts were separated along the same dimension as within-context (Figure 6F, left).

Having validated these constraints quantitatively, we concluded that population activity across gains formed two groups of isomorphic speed-dependent neural trajectories (Figure 6A). These results support our primary hypothesis that flexible control of speed based on gain context was established by a context-dependent persistent external input (Figure 3B).

**Figure 5.**
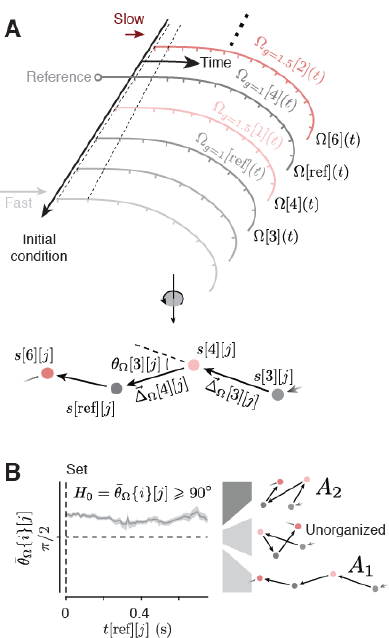
Neural trajectories across contexts do not form a single structure reflecting *t* _*p*_. (**A**) A schematic illustrating neural trajectories across the two contexts after Set. Top: The expected geometrical structure under *A* _*1*_. Neural trajectories for the gain of 1 (gray) and 1.5 (red) are organized along a single initial condition axis and ordered with respect to *t* _*p*_. Bottom: A rotation of the top showing neural trajectories with the time axis normal to the plane of illustration. If the neural trajectories were organized as such, then the angle between vectors connecting nearby points (e.g.,θΩ[3][*j*]) would be less than 90 (A *1*, Figure 3A). (B) Left: orientation of vectors connecting adjacent neural trajectories combined across the two contexts. Right: possible geometrical structures, including *A* _*1*_ (bottom), *A* _*2*_ (top), and unorganized (middle). 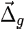 was larger 90 degrees for all *j* in the Set-Go interval, consistent with *A* _*2*_. Shaded regions represent 90% bootstrap confidence intervals.

**Figure 6.**
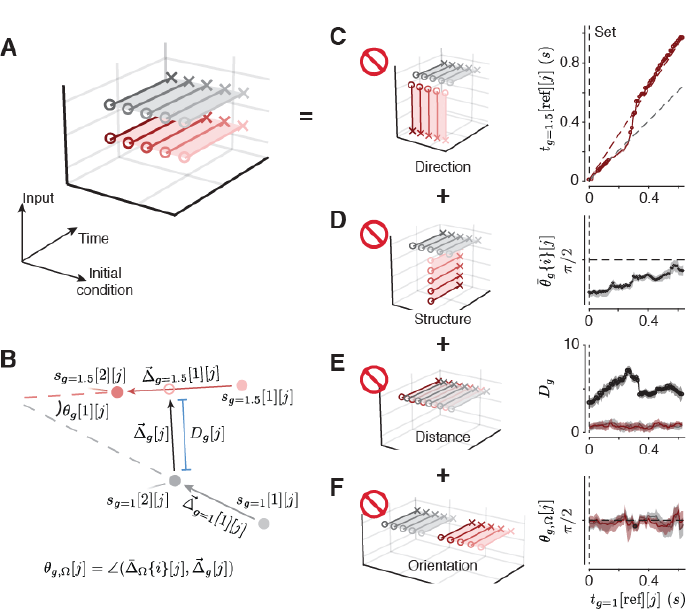
Neural trajectories comprise distinct but similar structures across gains. (**A**) A schematic showing the organization of neural trajectories in a subspace spanned by Input, Initial condition and Time if context were controlled by persistent external input. If DMFC were to receive a gain-dependent input, we would expect neural trajectories from Set to Go to be separated along an input subspace, generating two similar but separated *t* _*p*_-related structures for each context (*A* _*2*_, Figure 3B). We verified this geometrical structure by excluding alternative structures (interdictory circles indicate rejected alternatives). (B) An illustration of neural trajectories for g=1 (gray filled circle) and g=1.5 (red filled circle) with the time axis normal to the plane of illustration. Gray and red arrows show vectors connecting nearby points in each context independently (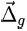 and 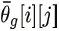). When the neural trajectories associated with the two gains are structured similarly, these vectors are aligned and the angle between them (θ^g^) is less than 90 deg. We used KiNeT to test this possibility (see Methods). (C) Left: Schematic illustrating a condition in which the time axis for trajectories in the two contexts (gray and red) are not aligned. Right: *t*^g^=1[ref]{*j*} increased monotonically with *t*^g^=1.5[ref]{*j*} indicating that the time axes across contexts were aligned. Values of above the unity line indicate that activity evolved at a slower speed in the g=1.5 context. The dashed gray line represents unity and the dashed red line represent expected values for *t*^g^=1.5[ref]{*j*} if speeds were scaled perfectly by a factor of 1.5. (D) Left: Schematic illustrating an example configuration in which Ω^g^=1{*i*}and Ω^g^=1.5{*i*} do not share the same *t* _*p*_-related structure. Right: 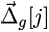 was significantly less than 90 degrees for all *j* indicating that the tp-structure was similar across the two contexts.(E). Left: Schematic illustrating a condition in which Ω^g^=1{*i*} and Ω^g^=1.5{*i*} are overlapping. Right: The minimum distance *D*^g^ across contexts (black line) was substantially larger than that found between subsets of trajectories within contexts (red and gray lines, see Methods) indicating the two sets of trajectories were not overlapping. (F) Left: Schematic illustrating a condition in which Ω^g^=1{*i*} Ω^g^=1.5{*i*} are separated along the same direction neural trajectories within each context were separated. Right: 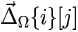 was orthogonal to 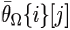 representing *t* _*p*_-related structure within each context (gray and red lines). In (C-E), shaded regions represent 90% bootstrap confidence intervals, and circles represent statistical significance (p < 0.05, bootstrap test, n = 100).

### RNN models recapitulate the predictions of inputs and initial conditions

The geometry and dynamics of DMFC responses were consistent with the hypothesis that behavioral flexibility in the RSG task relies on systematic adjustments of initial conditions and external inputs of a dynamical system. Motivated by recent advances in the use of recurrent neural networks (RNNs) as a tool for testing hypotheses about cortical dynamics (Mante et al. 2013; Hennequin et al. 2014; Sussillo et al. 2015; Chaisangmongkon et al. 2017; Wang et al. 2017), we investigated whether RNNs trained to perform the RSG task would establish similar geometrical structures and dynamics.

We focused on a generic class of RNNs comprised of synaptically coupled nonlinear units that receive nonspecific background activity (see Methods). First, we tested whether RNNs could perform the RSG task in a single gain context (*g* =1 or *g* =1.5 only). To do so, we created RNNs that received an additional input encoding Ready and Set as two brief pulses separated by *t s*. We trained these RNNs to generate a linear output function after Set that reached a threshold (Go) at at the desired production interval, *t t* = *gt s*. Analysis of successfully trained RNNs revealed that they, like DMFC, controlled *t p* by adjusting the speed of neural trajectories within a low-dimensional geometrical structure parameterized by initial conditions (Figure S4).

Next, we investigated RNNs trained to perform the RSG task across multiple gain values. Our primary aim was to verify the importance of a persistent gain-dependent input in establishing isomorphic geometrical structures similar to DMFC (Figure. 3, 6). To do so, we created RNNs with two different architectures, one in which the gain information was provided by the level of a persistent input, and another in which the gain information was provided by a transient pulse before the Ready cue. We refer to these networks as tonic-input RNNs and transient-input RNNs, respectively (Figure 7A). We used the tonic-input RNN as a direct test of whether a gain-dependent persistent input could emulate the geometrical structure of responses in DMFC, and the transient-input RNN to test whether such persistence was necessary.

Using PCA and KiNeT, we found that neural trajectories in the two networks were structured differently. In the tonic-input RNN, trajectories formed two isomorphic structures separated along the dimension associated with the gain-dependent persistent input (Figure 7B). In contrast, trajectories generated by the transient-input RNN were better described as coalescing towards a single structure parameterized by initial condition (Figure 7C). To verify these observations quantitatively, we evaluated the geometry of neural trajectories in the two RNN variants using the same analyses we performed on DMFC activity. In particular, we sorted trajectories with respect to *t p* across the two gain contexts (*g* =1 and *g* =1.5) and quantified the angle between vectors connecting nearest points (θΩ[*i*][*j*]). As noted in the analysis of DMFC, this angle is expected to be acute if trajectories form a single structure (*A 1*: 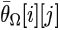 < 90 deg), and obtuse if trajectories form two gain-dependent structures (*A*2:θΩ[*i*][*j*] ≥90 degree). As predicted, the tonic-input solved the task by forming two isomorphic structures (*A*2) indicating that when a persistent gain-dependent input is present, RNNs rely on a solution with separate gain-dependent geometrical structures

We also compared the two RNNs in terms of the distance between trajectories across the two contexts using the same metric (*D*^g^) we used previously used for the analysis of DMFC (Figure 6E). The minimum distance between Ω ^*g*^ and Ω ^*g*^ at the time of Set was consistently smaller in the transient-input RNN compared to tonic-input RNN (Figure 7F,G). In some of the successfully trained transient networks, *D*^*g*^ was larger at the time of Set, but this distance consistently decayed from Set to Go. In contrast, in the tonic-input RNN,*D*^*g*^ remained large throughout the production epoch. We compared the two types of RNN quantitatively but comparing values of *D*^*g*^ in each RNN normalized by the distance between the trajectories that correspond to the shortest and longest *t*^*p*^ bin for the g= 1 context in the same RNN. In the tonic networks, the minimum normalized distance ranged between 0.4 and 1.6, which was nearly 10 times larger than the that observed in the transient networks (0.003 to 0.04). Additionally, trajectories in all transient networks gradually established a *t*_*t*_-related structure consistent with A 1. In contrast, trajectories in the tonic networks, like the DMFC data, were characterized by two separate *t*^*p*^-related structures, one for each gain context. These results provide an important theoretical confirmation of our original dynamical systems hypothesis that when gain information is provided as persistent input, the system establishes distinct and isomorphic gain-dependent sets of neural trajectories.

**Figure 7.**
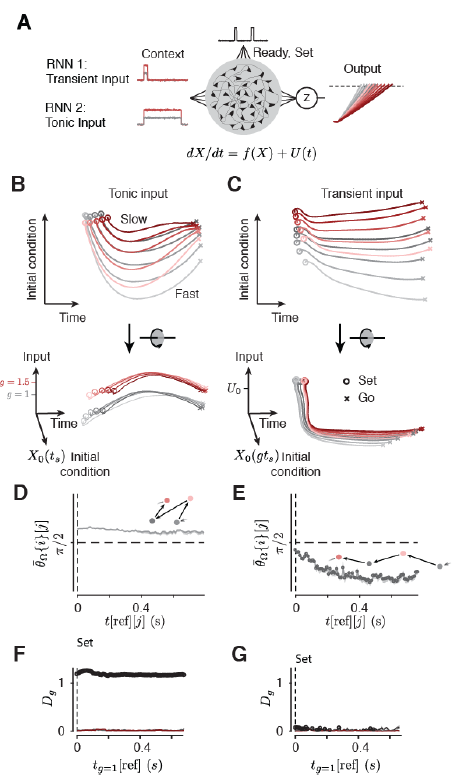
RNNs with tonic but not transient input captured the structure of activity in DMFC. (**A**) Schematic illustration of the recurrent neural networks (RNNs). The networks are provided with brief Ready and Set pulses separated in time by *t* _*s*_, after which the activity projected onto the output space by weighting function z must generate a ramp to a threshold (dashed line) at the context-dependent *t* _*t*_. Additionally, each network is provided with a context-dependent “input” which either terminates prior to Ready (“Transient input,” top), or persists throughout the trial (“Tonic input,” bottom) (B) Top: state-space projections of tonic-input RNN activity in the Set-Go epoch within the plane spanned by Initial condition (ordinate) and Time (abscissa). Within this plane of view, neural trajectories within each context are separated based on *t* _*p*_ but overlap with respect to gain. Bottom: Same neural trajectories shown in the top panel viewed within the plane spanned by Input (ordinate) and Time (abscissa). In this view, neural trajectories are separated by gain but overlap with respect to *t* _*p*_ within each gain. Results are shown with the same format as Figure 3. (C) Same as panel B for the transient-input RNN. Top: Trajectories, when viewed within the plane of Initial condition and Time, are organized with respect *t* _*p*_ across both gains. Bottom: when viewed within the plane of Input and Time, trajectories are highly overlapping irrespective of gain. (**D**) Analysis of direction in the tonic-input RNN with the same format as Figure 5B. 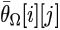 was larger than 90 deg for the entire the Set-Go epoch. This is consistent with a geometry in which the two gains form two separate sets of isomorphic neural trajectories (inset). (**E**) Same as panel D for the transient-input network for 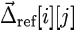 which was consistently less than 90 deg. This is consistent with a geometry in which neural trajectories are organized with respect to *t* _*p*_ regardless of the gain context (inset). (F,G) Trajectory separation across contexts for the tonic-input (F) and transient-input (G) networks with the same format as Figure 6E. *D* ^*g*^ was substantially larger through the Set-Go epoch in the tonic-input network (F). In (D-G), shaded regions represent 90% bootstrap confidence intervals, and circles represent statistical significance (p < 0.05, bootstrap test, n = 100).

## Discussion

Linking behavioral computations to neural mechanisms requires that the space of models we consider suitably match the computational demands of the behavior. In this study, we focused on the computations that enable the brain to exert precise and flexible control over movement initiation time (Wang et al. 2017). Because such temporal control depends on intrinsically dynamic patterns of neural activity, we employed a dynamical systems perspective to understand the underlying computational logic. An important feature of the dynamical systems view is that it obviates the need for the system to harbor an explicit representation of experimentally defined task-relevant variables (*t s, g*, and *t t*). Instead, neural signals that control behavior may be more appropriately characterized in terms of constraints imposed by latent dynamics that hold an implicit representation of task-relevant variables to control behavior. This viewpoint has a strong basis in current theories of motor control that posit an implicit representation of kinematic information in motor cortical activity during movements (Churchland et al. 2010; Churchland et al. 2012; Chaisangmongkon et al. 2017; Fetz 1992; Shenoy et al. 2013; Michaels et al. 2016). These theories cast movement control in terms of the function of an inverse model (Wolpert & Kawato 1998; Todorov & Jordan 2002; Sabes 2000) that inverts a desired endpoint to suitable control mechanisms during movement. We built upon this framework by evaluating the utility of dynamical systems theory in characterizing the control mechanisms the brain uses to produce a desired interval (*t t*) jointly specified by gain and *t s* (*t t* = *gt s*.

Results indicated that flexible control of behavior could be parsed in terms of systematic adjustments to initial conditions and external inputs of a dynamical system. Activity structure within each gain context indicated that the system‘s initial conditions controlled *t p* by parameterizing the speed of neural trajectories (Jazayeri &Shadlen 2015; Wang et al. 2017). The displacement of neural trajectories in the state space as a function of gain, and the lack of structural representation of *t p* across both gains suggested that DMFC received the gain information as a context-dependent tonic input. Following recent advances in using RNNs to generate and test hypotheses about dynamical systems (Mante et al. 2013; Rigotti et al. 2010; Hennequin et al. 2014; Rajan et al. 2016; Sussillo et al. 2015; Chaisangmongkon et al. 2017), we verified this interpretation by analyzing the behavior of different RNN models trained to perform the RSG task with either tonic or transient context-dependent inputs. Although both networks used initial conditions to set the speed of neural trajectories, only the tonic-input RNNs reliably established separate structures of neural trajectories across gains, similar to what we found in DMFC.

Although we do not know the constraints that led the brain to establish separate geometrical structures, we speculate about potential computational advantages associated with this particular solution. First and foremost, this may be a particularly robust solution; as the gain information was provided by a persistent visual cue, the brain could use this input as a reliable signal to modulate neural dynamics in RSG. This solution may also reflects animals’ learning strategy. We trained monkeys to perform the RSG tasks with two gain contexts. On the one extreme, animals could have treated these as completely different tasks leading to completely unrelated response structures for the two gains. On the other extreme, animals could have established a single parametric solution that would enable the animal to perform the two contexts as part of a single continuum (e.g., represent *t t*). DMFC responses, however, did not match either extreme. Instead, the system established what might be viewed as a modular solution comprised of two separate isomorphic structures. We take this as evidence that the brain sought similar solutions for the two contexts, but it did so while keeping the solutions separated in the state space. This strategy preserves a separable, unambiguous representation of gain and *t s* at the population level (Machens et al. 2010; Mante et al. 2013; Kobak et al. 2016) and provides the additional flexibility of parametric adjustments to the two parameters independently. Future extensions of our experimental paradigm to cases where context information is not present throughout the trial (e.g., internally inferred rules) might provide a more direct test of these possibilities.

Regardless of the learning strategies and constraints that shaped DMFC responses, our results highlight an important computational role for inputs that deviate from traditional views. We found that changing the level of a static input can be used to generalize an arbitrary stimulus response mapping in the RSG task to a new context. Similar inferences can be made from other recent studies that have evaluated the computational utility of inputs that encode task rules and behavioral contexts (Mante et al. 2013; Song et al. 2016; Chaisangmongkon et al. 2017). Extending this idea, it may be possible for the system to use multiple orthogonal input vectors to flexibly and rapidly switch between sensorimotor mappings along different dimensions. Together, these findings suggest that a key function of cortical inputs may be to flexibly reconfigure the intrinsic dynamics of cortical circuits by driving the system to different regions of the state space. This allows the same group of neurons to access a reservoir of latent dynamics needed to perform different task-relevant computations.

Our results raise a number of additional important questions. First, future work should identify the neurobiological substrate of the putative context-dependent input to DMFC in the RSG task, which may be among various cortical and subcortical areas (Lu et al. 1994; Bates & Goldman-Rakic 1993; Wang et al. 2005; Akkal et al. 2007; Wallis et al. 2001). The nature of the input is also unknown. In our RNN models, context information was provided by external drive, and was indistinguishable from recurrent inputs from the perspective of individual units. In cortex, reconfiguration of circuit dynamics may be achieved by either an external drive similar to the function of thalamic relay signals, or through targeted modulation of neural activity (Harris & Thiele 2011; Nadim & Bucher 2014). Second, while the signals recorded in this study were consistent with a prominent role for DMFC in RSG, other brain areas, such as the thalamus (Guo et al. 2017; Schmitt et al. 2017) and prefrontal cortex (Miller & Cohen 2001) are also likely to help maintain the observed dynamics. Third, although we assumed that recurrent interactions were fixed during our experiment, it is almost certain that synaptic plasticity plays a key role as the network learns to incorporate context-dependent inputs (Kleim et al. 1998; Pascual-Leone et al. 1995; Yang et al. 2014; Xu et al. 2009). Finally, the persistent separation of neural trajectories observed in DMFC allowed for a dynamical account which did not require invocation of “hidden” network states to explain timing behavior (Buonomano & Merzenich 1995; Karmarkar & Buonomano 2007; Murray & Escola 2017) or contextual control (Stokes et al. 2013). However, it is possible that factors not measured by extracellular recording (e.g., short-term synaptic plasticity) contribute to both contextual control and timing behavior in RSG and similar tasks. These open questions aside, our results provide a novel way to bridge the divide between neural activity and behavior by using the language of dynamical systems.

## Methods

All experimental procedures conformed to the guidelines of the National Institutes of Health and were approved by the Committee of Animal Care at the Massachusetts Institute of Technology. Two monkeys (*Macaca mulatta*), one female (C) and one male (J), were trained to perform the Ready, Set, Go (RSG) behavioral task. Monkeys were seated comfortably in a dark and quiet room. Stimuli and behavioral contingencies were controlled using MWorks (https://mworks.github.io/) on a 2012 Mac Pro computer. Visual stimuli were presented on a frontoparallel 23-inch Acer H236HL monitor at a resolution of 1920×1080 at a refresh rate of 60 Hz, and auditory stimuli were played from the computer‘s internal speaker. Eye positions were tracked with an infrared camera (Eyelink 1000; SR Research Ltd, Ontario, Canada) and sampled at 1 kHz.

### RSG Task

*Task contingencies.* Monkeys had to measure a sample interval, *t s t* whose relationship *to t s* was specified by a context-dependent gain parameter () which was set to either 1 (g=1 context) or 1.5 (g=1.5 context). On each trial, *t s* was drawn from a discrete uniform prior distribution (7 values, minimum = 500 ms, maximum = 1000 ms), and *gain* (*g*) was switched across blocks of trials (101 +/− 49 trials (mean +/− std)).

*Trial structure.* Each trial began with the presentation of a central fixation point (FP, circular, 0.5 deg diameter), a secondary context cue (CC, square, 0.5 deg width, 3–5 deg below FP), an open circle centered at FP (OC, radius 8–10 deg, line width 0.05 deg, gray) and three rectangular stimuli (2.0×0.5 deg, gray) placed 90 deg apart over the perimeter of OC with their long side oriented radially. FP was red for the g=1 context and purple for the g=1.5 context. CC was placed directly below FP in the g=1 context, and was shifted 0.5 deg rightward in the g=1.5 context. Two of the rectangular stimuli were presented only briefly and served as placeholders for the subsequent ‘Ready’ and ‘Set’ flashes. The third rectangle served as the saccadic target (‘Go’), which together with FP, CC, and OC remained visible throughout the trial. Ready was always positioned to the right or left of FP (3 o‘clock or 9 o‘clock position). Set was positioned 90 deg clockwise with respect to Ready and the saccadic target was placed opposite to Ready (Figure 1A).

Monkeys had to maintain their gaze within an electronic window around FP (2.5 and 5.5 deg window for C and J, respectively) or the trial was aborted. After a random delay (uniform hazard), first the Ready and then the Set cues were flashed (83 ms, white). The two flashes were accompanied by a short auditory cue (the “pop” system sound), and were separated by *t s*. The produced interval *t p* was defined as the interval between the onset of the Set cue and the time the eye position entered a 5-deg electronic window around the saccadic target. Following saccade, the response was deemed a “hit” if the error was smaller than a, and subsequently produce a target interval *t -* dependent threshold ∊*thresh*=α*tt*+β where α was between 0.2 and 0.25, β and was 25 ms. The exact choice of these parameters were not critical for performing the task or for the observed behavior; instead, they were chosen to maintain the animals motivated and willing to work for more trials per session. On hit trials, the target, animals received juice reward and FP turned green. The reward amount, as a fraction of maximum possible reward, decreased with increasing error according to ((∊*thresh*-∊)/,∊*thresh*)_1.5_, with a minimum fraction 0.1 (Figure 1B). Trials in which *t p* was more than 3.5 times the median absolute deviation (MAD) away from the mean were considered outliers and were excluded from further analyses.

As an initial analysis of whether monkeys learned the RSG task across gains, we fit linear regression models to the behavior separately for each gain:

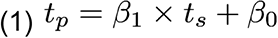

To quantify the difference in slopes between the two contexts. We also fit models with an interaction term across both contexts:

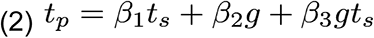

If the animals successfully learned to apply the gain, β should be positive.

We further applied a Bayesian observer model (Jazayeri & Shadlen 2015; Acerbi et al. 2012; Miyazaki et al. 2005; Jazayeri & Shadlen 2010), which captured the behavior in both contexts (Figure 1E). Full details of the model can be found in previous work (Jazayeri & Shadlen 2010; Jazayeri & Shadlen 2015). Briefly, we assumed that both measurement and production of time intervals are noisy. Measurement and production noise were modeled as zero-mean Gaussian with standard deviation proportional to the base interval (Rakitin et al. 1998), with constant of proportionality of of *w m* and *w p*, respectively. A Bayesian model observer produced*p* after deriving an optimal estimate of *t t* from the mean of the posterior. To account for the possibility that the mental operation of mapping *t s* to *t t* according to the gain factor might be noisier in the g=1.5 context than in*p* to vary across contexts. *t t t*

### Recording

We recorded neural activity in dorsomedial frontal cortex (DMFC) with 24-channel linear probes (Plexon, inc.). Recording locations were selected according to stereotaxic coordinates and the existence of task-relevant modulation of neural activity. In monkey C, recordings were made between 3.5 mm to 7 mm lateral of the midline and 1.5 mm posterior to 4.5 mm anterior of the genu of the arcuate sulcus. In monkey J, we recorded from between 3 mm to 4.5 mm lateral of the midline and 0.75 mm to 5 mm anterior of the genu of the arcuate sulcus. Data were recorded and stored using a Cerebus Neural Signal Processor (NSP; Blackrock Microsystems). Preliminary spike sorting was performed online using the Blackrock NSP, followed by offline sorting using the Phy spike sorting software package using the spikedetekt, klusta, and kilosort algorithms (Rossant et al. 2016; Pachitariu et al. 2016). Sorted spikes were then analyzed using custom code in MATLAB (The MathWorks Inc.).

### Analysis of DMFC data

Average firing rates of individual neurons were estimated using a 150 ms smoothing filter applied to spike counts in 1 ms time bins. We used PCA to visualize and analyze activity patterns across the population of neurons across animals. PCA was applied after a soft normalization: spike counts measured in 10 ms bins were divided by the square root of the maximum spike count across all bins and conditions. The normalization was implemented to minimize the possibility high firing rate neurons dominating the analysis.

When binning data according to increasing values of *t p*, we ensured that all bins had equal number of trials, independently for each session. To average firing rates across trials within a group, we truncated trials to the median *t p*, and averaged firing rates with attrition. Analyses of neural data were applied to all 10 sessions across both monkeys. For analyses, we included neurons for which at least 15 trials were recorded in each condition and which had a minimum unsmoothed modulation depth of 15 spikes per second. We did not separately analyze trials immediately following context switches due to the low number of context switches per session (mean = 6.8 switches).

For visualization of neural trajectories in state space, we identified dimensions along which responses were maximally separated with respect to context (“gain axis,” Figure 2B,C, “initial condition,” Figure 3C) and *t p* (“interval axis,” Figure 2B,C, and “initial condition,” Figure 3C). We first calculated the context component by projecting data onto the vector defined by the difference between neural activity averaged over time and *t p* for each context. This component of the activity was then subtracted away from the full activity. For the Ready-Set epoch, we then performed PCA (PCs 1 and 2, Figure 2B) on the data with the context component removed.For the Set-Go epoch, we calculated the *t p* component by projecting data onto the vector defined by the difference between the activity associated with longest and shortest values *t p*, averaged across time and) on the data with the context and *t p* components context. We then performed PCA (PC 1, Figure. 2C and 3C removed.

### Kinematic analysis of neural trajectories (KiNeT)

We developed KiNeT to compare the geometry, relative speed and relative position along a group of neural trajectories that have an orderly organization and change smoothly with time. To describe KiNeT rigorously, we developed the following symbolic notations. Square and curly brackets refer to individual items and groups of items, respectively.

The algorithm for applying KiNeT can be broken down into the following steps: 1) Choose a Euclidean coordinate system to analyze the neural trajectories. We chose the first 10 PCs in the Set-Go epoch, which captured 89% of the variance in the data. 2) Designate one trajectory as reference, Ω[ref]. We used the trajectory associated with the middle *t p* bin as reference. 3) On each of the non-reference trajectories Ω[ref](*i*≠ref), find *s*[*i*{*j* with minimum Euclidean distance to *s*[ref]{*j*} and their associated times *t*[*i*]{*j*}according to the following equations:

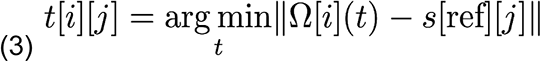

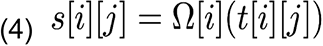

Organization of trajectories in state space: The distances *D*[*i*]{*j*}were used to characterize positions in neural state space of each Ω[*i*] relative to Ω[ref]. The magnitude of *D*[*i*][*j*] was defined as the norm of the vector connecting *s*[*i*][*j*]to *s*[ref][*j*], which we refer to as 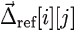. The sign of *D*[*i*][*j*]was defined as follows: for the trajectory Ω[1] associated with the shortest *ts* or *tp*, and Ω[*N*] associated with the longest, *D*[*i*][*j*] was defined to be negative and positive, respectively. For all other trajectories, *D*[*i*][*j*] was positive if the angle between 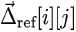 and 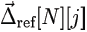 was smaller than the angle between 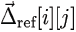 and 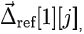, and negative otherwise.

Analysis of neural trajectories across contexts: We analyzed the geometry across gains in three ways. First, we analyzed the relationships between the two sets of trajectories. This required aligning the activity between the two contexts in time. To do this, we started with the aligned times *t*{*i*}{*j*} found within each context, and using successive groups of neural states in the g=1 context indexed by *tg*=1[ref]{*j*}, found the reference time *tg*=1.5[ref]{*j*} in the g=1.5 context for which the mean distances between neural states in paired trajectories (i.e. the first *t p* bins of both gains, second *t p* bins, etc.) were smallest. This resulted in an array of times from *tg*=1.5[ref]{*j*}, indexed by *tg*=1[ref]{*j*}, such that the trajectories across gains were aligned in time for subsequent analyses (Figure 6C). The second way that we analyzed geometry across gains was to collect trajectories across both gains, order according to trajectory duration, and run the standard KiNeT procedure. Finally, we measured the distance between the structures using the across-context time alignment. For successive, we measured the minimum distance between line segments connecting consecutive trajectories within each context. For five *t p* bins, this meant four line segments for each context, and 4 2 =16 distances. We chose the minimum of these distance values as the value of between the two structures. As a point of comparison, we generated set of “null” distances by splitting trajectories from each context into odd- and even-numbered trajectories and calculating the minimum distance between the sets of connecting line segments Figure 6E).

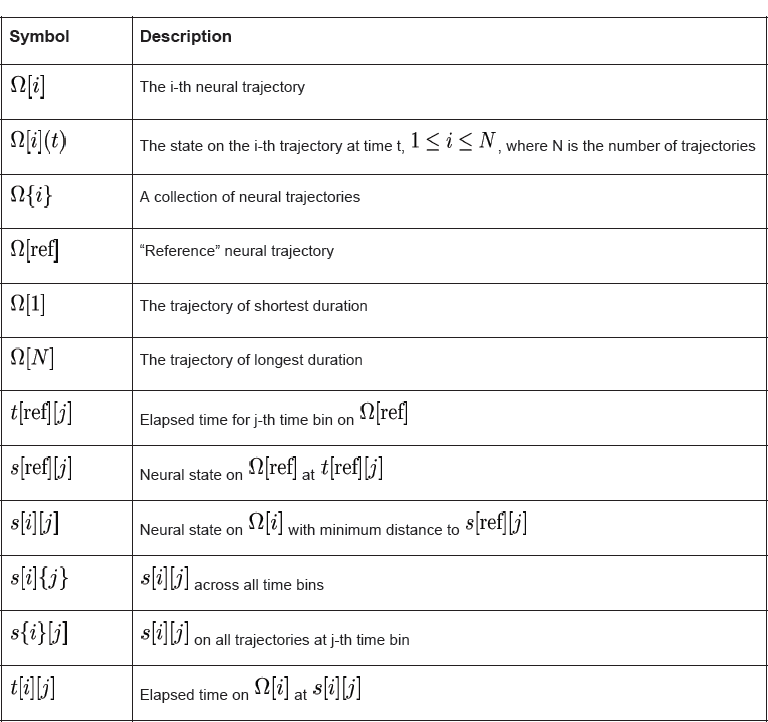

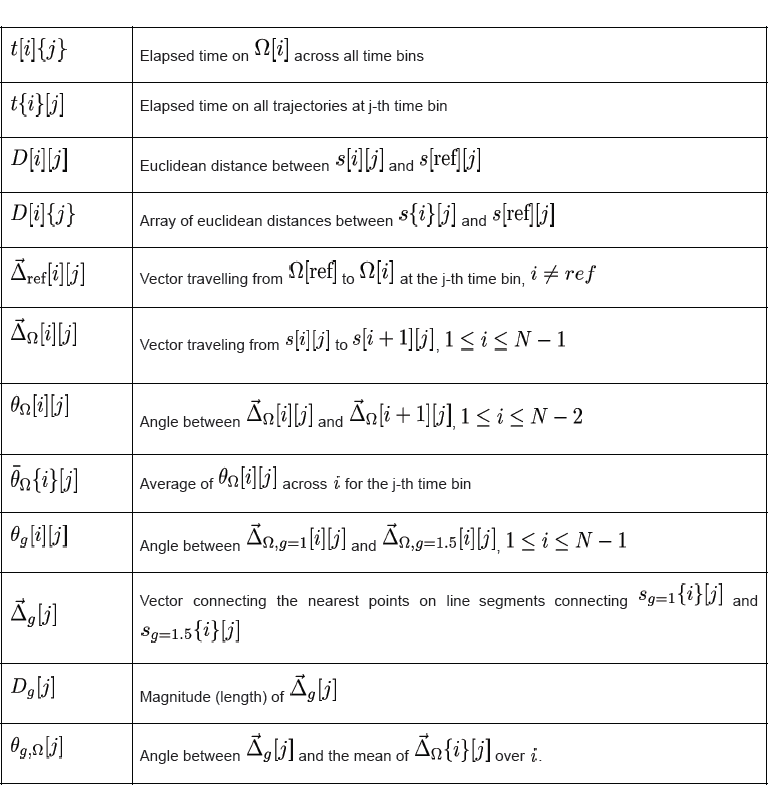

Statistics: Confidence intervals for KiNeT performed on trajectories binned according to *t p* were computed by a bootstrapping procedure, randomly selecting trials with replacement 100 times. To test for statistical significance of metrics generated through the KiNeT procedure, we used bootstrap tests, where p was the fraction of bootstrap iterations for which the metric was consistent with the null hypothesis. Unless otherwise stated, significance of a measure for individual time points was set to p < 0.05. The results of KiNeT applied to neural data from individual monkeys produced similar results, and were similar for different methods of data smoothing.

### Recurrent neural network

We constructed a firing rate recurrent neural network (RNN) model with *N* = 200 nonlinear units. The network dynamics were governed by the following differential equation:

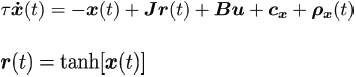

***x***(*t*) is a vector containing the activity of all units. and ***r***(*t*) represents the firing rates of those units by transforming *x* through a *tanh* nonlinearity. Time *t* was sampled every millisecond for a duration *T* of = 3300 ms. The time constant of decay for each unit was set to *t*=10ms. The unit activations also contain an offset ***c*** *x* and white noise ***p****x*(*t*) at each time step with standard deviation in the range [0.01-0.015]. The matrix ***j*** represents recurrent connections in the network. The network received multi-dimensional input ***u*** through synaptic weights ***b***=[***b****c*,***b****s*]. The input ***u*** was comprised of a gain-dependent context *uc* (*t*) and an input *us* (*t*) that provided Ready and Set pulses.In *us* (*t*) Ready and Set were encoded as 20 ms pulses with a magnitude of 0.4 that were separated by time *t*_*s*_.

Two classes of networks were trained to perform the RSG task with multiple gains. In the tonic-input RNNs, the gain-dependent *uc* (*t*) input was set to a fixed offset for the entire duration of the trial. In contrast, in the transient-input RNNs, *us* (*t*) was active transiently for 440 ms and was terminated 50-130 ms before the onset of the Ready pulse. The amplitude of *us* (*t*) was set to 0.3 for g=1 and 0.4 for g=1.5. The transient network received an additional gain-independent persistent input of magnitude 0.4, similar to the tonic networks. Both types of networks produced a one-dimensional output *z*(*t*) through summation of units with weights ***w****o* and a bias term *cz*.

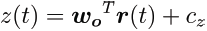

### Network Training

Prior to training, model parameters (θ), which comprised ***J***,***B***,***W****o*, ***c****z* and were initialized. Initial values of matrix ***J*** were drawn from a normal distribution with zero mean and variance 1/ *N* (Rajan & Abbott 2006). Synaptic weights***B***=[***b****c*,***b****s* and the initial state vector ***x***(0) and unit biases***c****x* were initialized to random values drawn from a uniform distribution with range [-1,1]. The output weights,***w****o* and bias ***c****z*, were initialized to zero. During training, model parameters were optimized by truncated Newton methods using backpropagation-through-time (Werbos 1990) by minimizing a squared loss function between the network output *zi*(*t*) and a target function *fi*(*t*), as defined by:

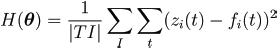

Here *i* indexes different trials in a training set (*I* = different gains (*g*) x intervals (*ts*) x repetitions (*r*)). The target function *fi*(*t*) was only defined in the Set-Go epoch (the output of the network was not constrained during the Ready-Set epoch). The value *fi*(*t*) of was zero during the Set pulse. After Set, the target function was governed by two parameters that could be adjusted to make *fi*(*t*) nonlinear, scaling, non-scaling or approximately-linear:

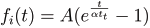

For the networks reported, *fi*(*t*) was an approximately-linear ramp function parametrized by *A*= 3 and α= 2.8. Variable *tt* represents the transformed interval for a given*ts* and gain *g*. Solutions were robust with respect to the parametric variations of the target function (e.g., nonlinear and non-scaling target functions). In trained networks, the production time, *t* _*p*_ was defined as the time between the Set pulse and when the output ramped to a fixed threshold (*z*_*i*_=1).

During training, we employed two strategies to obtain robust solutions. First, we trained the networks to flexibly switch between three gain contexts, the two original values (*g* =1 and *g* =1.5) and an additional intermediate value of *g* =1.25 for which the amplitude of *u*_c_(*t*) was set to 0.35. However, the behavior of networks trained with the two original gains were qualitatively similar. Second, we set ***P***(***t***) to zero, and instead, the context-dependent input,*u*_c_(*t*) received white noise with standard deviation of 0.005, per unit time (δ*t*=0).

## Supplement

### Go-aligned KiNeT

**Figure S1.**
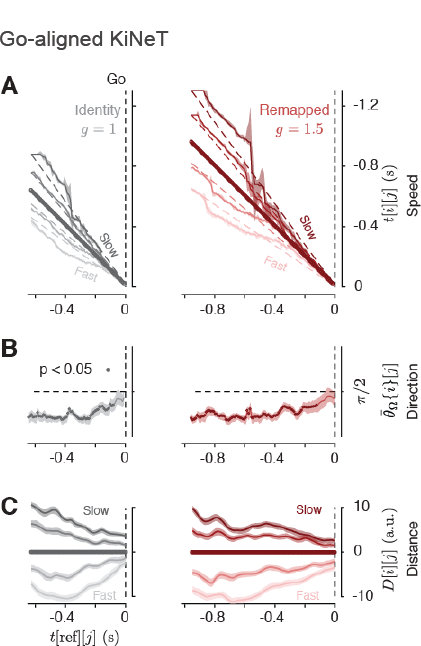
“Go“-aligned KiNeT, related to Figure 4. Applying KiNeT to neural trajectories aligned to the Set cue resulted *t*[*i*][*j*] in which diverged from *t*[ref] to scale with trajectory length in a manner consistent with neural speed control as a means to produce different *t* _*p*_. To rule out the possibility that this temporal scaling of trajectories was an artifact of temporal smearing of PSTHs near the time of Go caused by averaging trials of different lengths, we applied KiNeT to data aligned to Go (saccade). (A). Aligned times (speed) across both contexts. As in the Set-aligned analysis,*t*[*i*][*j*] for shorter Ω[*i*] diverged to shorter values, while *t*[*i*][*j*] for longer Ω[*i*] diverged towards longer values as *t*[ref] (here time before Go) increased. In contrast to the lack of temporal scaling proximal to the Set cue, *t*[*i*][*j*] were ordered according to *t* _*p*_ leading all the way up to the Go cue. Circles on the *t*[ref] line indicate ***j*** for which the ordering of *t*[*i*][*j*] was significantly correlated with the *t* _*p*_ bin (bootstrap test, r > 0.1, p < 0.05, n = 100). (B,C) *t* _*p*_-related structure of Ω{*i*} (B). Analysis of direction. As in the Set-aligned KiNeT, 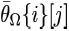 (bar signifies mean over the index in curly brackets) was significantly smaller than 90 degrees for the majority of the Set-Go interval (bootstrap test, 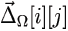 < 90, p < 0.05, n = 100) indicating that 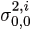 were similar, across Ω[*i*]. (C) Analysis of distance. Euclidean distance to Ω[ref]. Trajectories were ordered in neural space according to *D*[*i*][*j*], where Ω[*i*] with *t* _*p*_ with more similar to the middle *t* _*p*_ bin to being located closer to Ω[ref]. Significance tested by counting the number of times in which *D*[*i*][*j*] was not ordered according to *t* _*p*_ bin in bootstrap samples for each *j* (p < 0.05, n = 100).

### Rotation of trajectories through time

**Figure S2.**
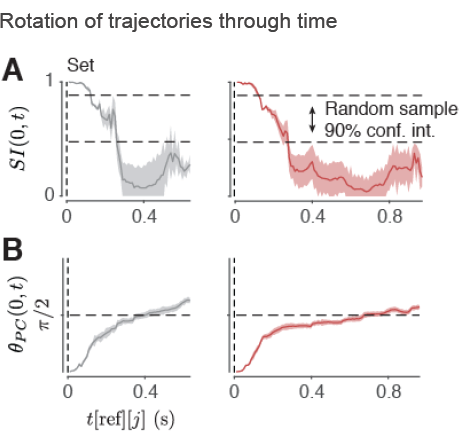
Rotation of trajectories through time, related to **Figure 4**. We estimated the degree to which the principal axes (PC directions) associated with nearest states along the five trajectories,*s*{*i*}[*j*], changed with time relative to t=0 using two metrics: a similarity index (*SI*(0,*t*) that measures the variance explained by PCs at time t and t = 0 (see below for full description), and a rotation index (θ_*PC1*_(0,*t*)) measuring the angle of the first PC (PC 1) in the state space at time t compared to t = 0. (A) ***SI***(0,*t*). This index varies between 0 and 1 with 1 signifying matching PCs and 0 signifying orthogonal PCs. The gradual change in (***SI***(0,*t*) away from 1 and toward 0 indicated that Ω{*i*} gradually changed orientation with time. Shaded area represents 90% bootstrap confidence intervals (n = 100). Dashed lines represent the 90% confidence intervals for the similarity of two sets of *s*{*i*}[*j*] drawn randomly from a multivariate Gaussian distribution with covariance matched to the data. (***SI***(0,*t*), captures the extent to which the orientation of *s*{*i*}[*j*] in state space changes *D*[*i*][*j*] with time and is therefore sensitive to both rotations and scaling transformations. (B)(θ_*PC1*_(0,*t*)). The gradual change in (θ_*PC1*_(0,*t*)) away from 0 toward 90 deg indicates that trajectories underwent rotations through state space from Set to Go. Unlike ***SI***(0,*t*) that is sensitive to both rotations and scaling transformations,(θ_*PC1*_(0,*t*)) is only sensitive to rotations. These data-driven observations motivated the use of KiNeT for analyzing neural trajectories throughout the paper.

Similarity Index: The similarity index, adapted from (Garcia 2012), was calculated using the following procedure: 1) Select two datasets, one for neural activity patterns at the time of Set (*t=0*), denoted by, and one at time *t* after Set, denoted by ***r***_*t*_. 2) Calculate the principal component coefficients for each dataset. 3)Project the points of each dataset onto their own and the others’ principal coefficients, creating four sets of principal component scores. 4) Calculate the fraction of variance explained by each principal component in each of the four sets of scores. 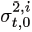 is the fraction of variance in ***r***_*t*_ explained by principal component *i* of ***r***_*o*_, 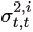 is the fraction of variance of ***r***_*t*_ explained by principal component *i* of ***r***_*o*_ 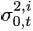 is the fraction of variance in explained by principal component *i* of ***r***_*t*_, and 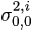 is the fraction of variance in ***r***_*o*_ explained by principal component *i* of ***r***_*t*_. 5) For each component of each dataset, calculate the difference between (1) the fraction of variance explained by that component for its own dataset (e.g.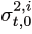 and (2) the fraction explained by that same component for the other dataset (e.g. 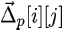). 6). Sum and normalize the calculated differences. This can be written as follows:

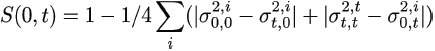

The similarity index is 0 when the associated covariance matrix of one dataset lies in the nullspace of the other, and 1 when the covariance matrices are identical.

In order to interpret the values of similarity index in the DMFC dataset, we compared similarity index for two surrogate datasets that matched the statistics of DMFC activity. Each dataset was constructed by drawing five samples (the number of *t p* bins) from a ten-dimensional Gaussian distribution (the number of principal components) with a diagonal covariance matrix constructed using the eigenvalues of the covariance matrix of the DMFC data. We calculated the similarity index for 1000 pairs of surrogate data (i.e., null distribution), and used the 5th and 95th percentiles to generate 90% confidence intervals. With this procedure, a similarity index above the 90% confidence interval was considered more “similar” than expected by chance, whereas a similarity index below the 90% confidence interval was considered dissimilar.

### Variability in neural trajectories systematically predicted behavioral variability

**Figure S3.**
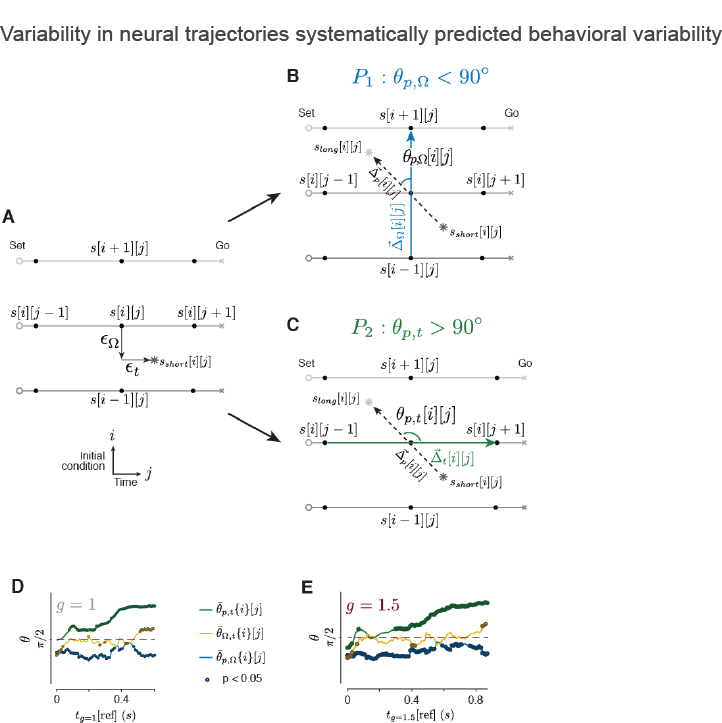
Relating neural variability to behavioral variability, related to Figure 4. (A). Schematic showing three neural trajectories between Set (circle) and Go (cross) associated with three different *t s* values. Neural states,*s*[*i*[*j*] indexed by trajectory (*i*), which is specified by initial condition, and elapsed time (*j*). Noise may cause neural states to deviate from mean trajectories. We reasoned that deviations across and along trajectories may cause systematic biases in *t p*. *s*_*short*_[*i*][*j*] (light star) shows an example in which noise moves the state in the direction of shorter *t s* (toward *s*[*i*-1][*j*]) and in the direction of the Go state (*s*[*i*][*j*+1]) by vectors ∊_Ω_ and ∊_t_, respectively. Both deviations should lead to shorter *t p*. (B) Prediction 1 (*P 1*): deviations off of one trajectory toward a trajectory associated with larger *t s* should lead to larger *t p*, and vice versa. To test*P 1*, we divided trials for each *t s* into two bins. One bin contained all trials in which *t p* was shorter than median *t p* and the other, all trials in which *t* |*p* was longer than median *t p*. We computed neural trajectories for the short and long *t p* bins, and denoted the corresponding states by *s*_*short*_[*i*][*j*] and *s*_*long*_[*i*][*j*](dark star), respectively. If*P 1* is correct, then the geometric relationship between *s*_*short*_[*i*][*j*] and *s*_*long*_[*i*][*j*]should be similar to that between *s*[*i*-1][*j*] (shorter *t s*) and *s*[*i*][*j*+1])(longer *t s*. Therefore the vector pointing from *s*_*short*_{*i*][*j*] to *s*_*long*_{*i*][*j*](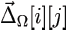, dashed arrow) and the vector pointing from *s*[*i*-1][*j*] to*s*[*i*][*j*+1], 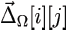 blue arrow) should be aligned, and the angle between them, denoted by θ_p_,Ω[*i*][*j*] should be acute. See below description for calculation of 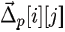 for shortest and longest *t s* along.(C) Prediction 2 (*P 2*): deviations ∊_*t*_ trajectories should influence the time it takes for activity to reach the Go state and should therefore influence *t p* (Afshar et al. 2011; Michaels et al. 2015). If *P 2* is correct, then *s*_*short*_[*i*][*j*] should be ahead of *s*_*long*_[*i*][*j*]. Therefore, 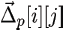 should point backwards in time, and the angle between 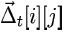 and 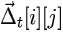 that connects *s*[*i*-1][*j*] to*s*[*i*][*j*+1], denoted by θ_*p*_,_*t*_[*i*][*j*] should be obtuse. See below description for calculation of 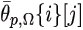 for first and last time points.

(D,E) Testing *P 1* and *P 2* for the *g* =1 (D) and *g* =1.5 (E) contexts. Consistent with *P 1*, average θ_p_,Ω{*j*}[*j*] (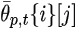, blue), were less than 90 deg from Set to Go indicating that *t p* was larger (smaller) when neural states deviated toward a trajectory associated with a larger (smaller) *t s*. Importantly, the systematic relationship between *t p* and neural activity was already present at the time of the Set, indicating that *t p* was influenced byvariability during the Ready-Set measurement epoch. Consistent with *P 2*, 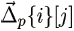 (green) was greater than 90 deg, indicating that *t p* was larger (smaller) when speed along the neural trajectory was slower (faster). The angle between 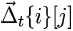 and 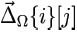 was initially close to 90 deg consistent with the observation that trajectories evolved at similar speeds early in the Set-Go epoch (Figure 4B).

We also measured the angle between 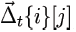 and 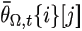, denoted by 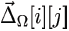 (yellow). This angle was not significantly different than expected by chance (90 deg) for most time points. We determined when (at what) an angle was significantly different from 90 deg (p < 0.05) by comparing angles to the corresponding null distribution derived from 100 random shuffles with respect to *t p*. Angles that were significantly different from 90 deg are shown by darker circles. Because the comparison of *t s* - vs. *t p*-related structure (Figure S3) required grouping trials into substantially more bins than the other analyses (14 vs. 7 or 5), we reduced the minimum number of trials required to 10 for this analysis (273 units; 95 from monkey C and 178 from monkey J). We did not find that the results of any of the analyses were dependent on the specific threshold chosen, and results were similar in individual subjects.

Calculation of 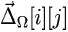 for shortest and longest *t s*: Because there was a finite number of *t s* values, we could not compute 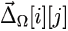 for *i*=1 and *i*=*N*_*i*_(*s*[*i*-1][*i*] was not defined for *i*=1 and*s*[*i*+1][*i*] was not defined for *i*=*N*_*i*_). Therefore, for the shortest t *s*, we changed to*s*[2][*j*]-*s*[1][*j*] (instead of)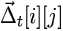 to*s*[2][*j*]-*s*[0][*j*], and for the longest *t s* [*N*_*i*_]-*s*[*N*_*i*_-1][*j*], to (instead of *s*[*N*_*i*_+1][*j*]-*s*[*N*_*i*_-1][*j*]).

Calculation of 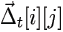 for the earliest and latest times: Because there was a finite number of time points, we could not compute 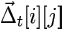 for *j*=1 and *j*=*N*_*j*_(*s*[*i*][*j*] was not defined for *j*=*N*_*j*_). Therefore, for the first time point, we changed 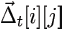 to *s*[*i*][2]-*s*[*i*[1](instead of *s*[*i*][2]-*s*[*i*][0]), and for the last time point, to *s*[*i*][*N*]-*s*{*i*][*N*_*j*_-1] (instead of *s*[*N*_*j*_+1][*j*]-*s*[*N*_*j*_-1][*j*]).

### Analysis of the recurrent neural networks

**Figure S4.**
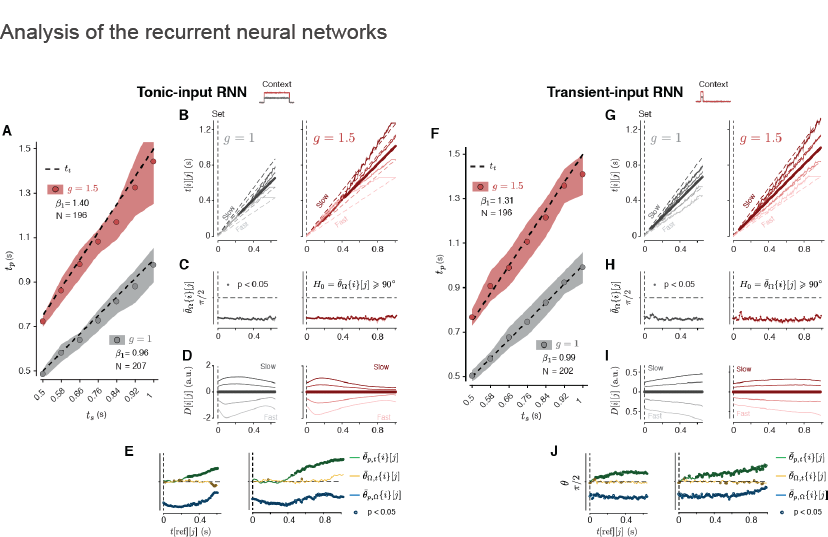
Analysis of the recurrent neural networks (RNNs), related to Figure 4 and Figure 7. (A-E) Tonic-input RNN. (A) “Behavior“; same format as in Figure 1E. The networks successfully learned the task as evidenced by positive regression slopes (, larger for *g* = 1.5 context) and a significant positive interaction between *t* _*s*_ and *g* (p << 0.001). For each network, we simulated 30 trials per *t* _*s*_ and *g*, removing outliers in which *t* _*p*_ was more than 3.5 times the median absolute deviation (MAD) away from the mean. (B-D) Organization of neural trajectories within each context; same format as Figure 4B-D. KiNeT analysis verified that the organization of neural trajectories in the tonic-input RNN matched the organization observed in DMFC (compare to Figure 4B-D). (E) Relating unit variability to behavioral variability; same format as in Figure S3. (F-J) Same analyses as in A-E for the transient-input RNN.

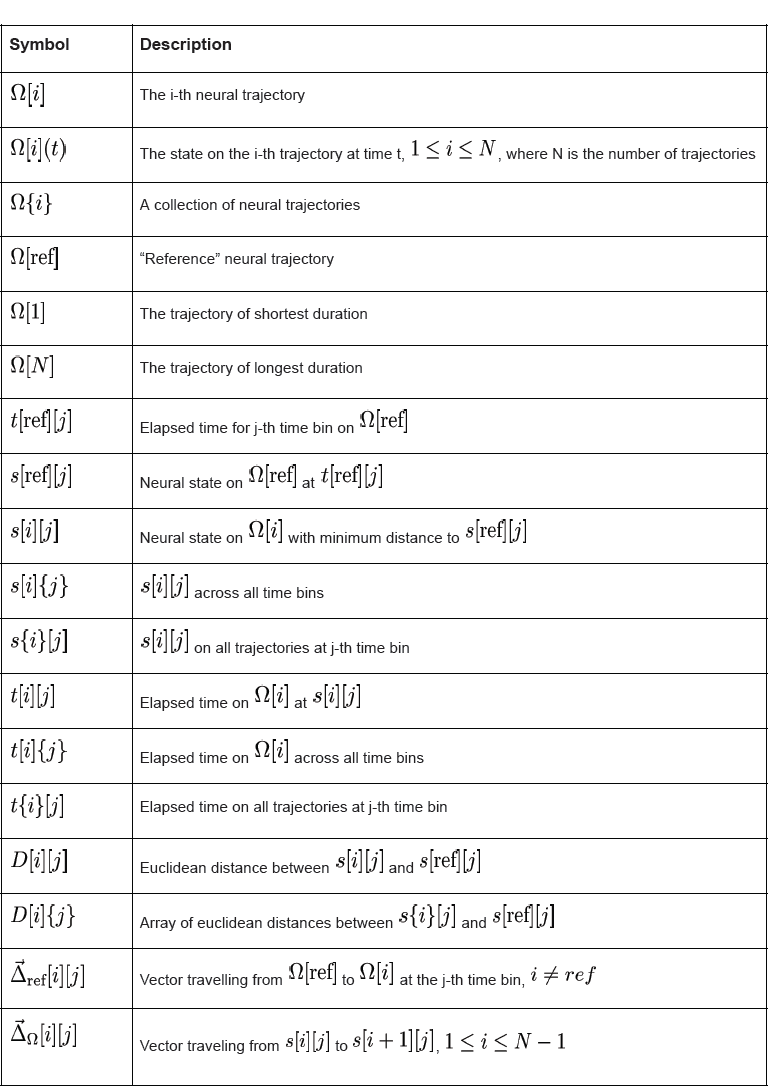

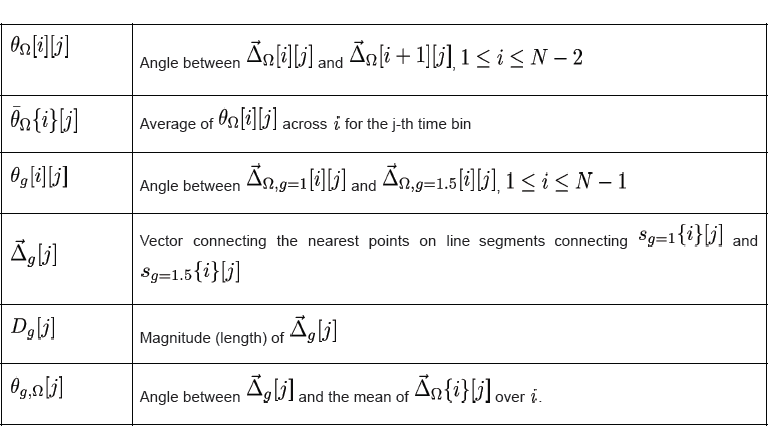

## Acknowledgements

We thank S.W. Egger, H. Sohn, and V. Parks for their helpful suggestions on the manuscript. D.N. is supported by the Rubicon grant (2015/446-14-008) from the Netherlands organization for scientific research (NWO). M.J. is supported by NIH (NINDS-NS078127), the Sloan Foundation, the Klingenstein Foundation, the Simons Foundation, the McKnight Foundation, the Center for Sensorimotor Neural Engineering, and the McGovern Institute.

## Author contributions

E.D.R. and M.J. designed the main experimental paradigm. E.D.R. and E.A.H. trained the animals. E.D.R. collected neural data from both animals. E.D.R. developed KiNeT. E.D.R. performed all analyses. D.N. developed the recurrent neural network models. E.D.R. and M.J. interpreted the results and wrote the paper.

